# A specific computational role for early-life unpredictability, and not lifelong traumatic experience, in decision-making under uncertainty

**DOI:** 10.64898/2026.06.01.729407

**Authors:** Yifan Zhang, Yifei Chen, X. Roger Chen, Nora C. Harhen, Laura A. Glynn, Elysia P. Davis, Tallie Z. Baram, Victoria B. Risbrough, Daniel A. Stout, Aaron M. Bornstein

## Abstract

It has been shown that early-life adversity (ELA) shapes how individuals learn, remember, and make decisions, yet the precise computations altered by these experiences remain unclear. Here, we combine a structured foraging task with computational modeling to test a recently developed theory for how a particular kind of ELA, early-life unpredictability (ELU), specifically influences choice under uncertainty. Adult participants (N=297) performed a sequential foraging task requiring continuous trade-offs between exploiting depleting resources and exploring alternatives. Subsets also completed assessments of early-life unpredictability (QUIC) and for trauma symptoms arising from lifelong stressors (PCL). We fit participants’ behavior with a Bayesian learning-and-planning model in which uncertainty modulates the valuation of leaving a current resource patch. Critically, the subjective influence of local uncertainty was allowed to vary freely between participants. Consistent with theoretical proposals, computational model fits and mediation analyses revealed a robust indirect pathway: ELU predicted increased discounting in the face of uncertainty, which in turn predicted greater overharvesting. This pattern was consistent across environmental conditions, indicating that early-life unpredictability primarily influences behavior through a general influence on uncertainty processing. Importantly, although PCL scores were also correlated with uncertainty adaptation, these effects were fully accounted for by shared variance with ELU, offering a clear dissociation between developmental unpredictability and lifelong traumatic experience. Together, our results show that early-life unpredictability causes long-lasting changes in decision-making by amplifying the subjective experience of uncertainty.

## Introduction

Early-life unpredictability (ELU) has been proposed as a critical developmental factor influencing learning and decision-making (Glynn et al., 2019; Birnie and Baram, 2022; Decker et al., 2025; Rachum et al., 2025). Exposure to unstable or unpredictable environments during development may alter how individuals represent and respond to uncertainty, potentially leading to persistent differences in valuation and action selection. Prior work has linked ELU to risky behaviors (Spadoni et al., 2022) and differences in affective processing (Glynn et al., 2019). A recent theoretical framework suggests that these behaviors are consistent with amplified uncertainty signals arising from adaptations during a critical period of high plasticity (Harhen and Bornstein, 2024). Here, we test this proposed specific computational role directly.

One promising framework for addressing the specific computational role in question is patch foraging, a decision behavior widely understood to engage naturalistic, evolutionarily-conserved mechanisms (Blanchard and Hayden, 2015). In canonical models (Charnov, 1976), decisions to leave a patch depend on a comparison between the expected value of staying and the average reward rate of the environment. When environmental structure is uncertain, agents must infer latent dynamics to estimate local and global patch richness, and adjust their decision policies accordingly (Harhen and Bornstein, 2023; Harhen et al., 2026; Chen et al., 2026). This suggests that individual differences in how uncertainty is estimated could lead to systematic variation in foraging behavior.

Later-life and acute stresses also affect uncertainty processing (De Berker et al., 2016; Morgado et al., 2015; Verfaellie et al., 2025). Lifelong experiences of trauma could both affect the content of self-report and also yield similar mental health outcomes as do early-life experiences. An open question for the field is whether the impact of these separate events on cognition can be dissected in adult behavior. A related, persistent concern when evaluating the influence of early-life experience on adult behavior is that measurement instruments used to evaluate early-life and other experience–typically, self-report questionnaires (Glynn et al., 2019)–could potentially be influenced by mental state at the time of elicitation. Despite extensive evidence of the validity of these instruments (Glynn et al., 2019; Spadoni et al., 2022), the question of interactive influences remains. For instance, a recent study found that depressive symptoms at the time of self-report influenced the recall of adverse childhood experiences (Zhang et al., 2026). Here, we address these concerns by asking participants to complete survey measures about both early-life and lifelong experience, and in a different session than the experimental task.

More specifically, we combine a sequential foraging task with computational modeling to test the hypothesis that ELU modulates uncertainty-dependent weighting. Participants completed a foraging task with patches of varying rewardingness. Rewards within each patch depleted over time, compelling participants to choose whether to continue harvesting the declining rewards or incur a travel cost and the possibility of transiting to a less (or more) rewarding patch. They also completed assessments of early-life unpredictability (QUIC) and trauma-related symptom severity (PCL). We fit participants’ choices with an Bayesian learning-and-adaptive discounting model (Harhen and Bornstein, 2023; Harhen et al., 2026; Chen et al., 2026) in which trial-by-trial uncertainty about the participants’ position in the state space adaptively modulates the value of leaving, captured by an uncertainty-dependent discount factor.

Consistent with prior theoretical work on the consequences of ELU for neural coding (Harhen and Bornstein, 2024), we hypothesized that individuals with greater ELU would exhibit increased sensitivity to uncertainty, which normative models (Jiang et al., 2015) suggest should be reflected in modulation of decision values by local uncertainty, specifically via a reduction in planning horizon. Furthermore, we tested whether this relationship is specific to ELU or can also be explained by lifelong trauma-related symptom severity. By linking early life experience to computational parameters governing decision-making, via normative models that formalize rational responses to life experience and environmental conditions, this work aims to provide a mechanistic account of how developmental environments shape adaptive behavior in uncertain and dynamic contexts.

## Materials and Methods

### Participants

297 participants were recruited from Amazon Mechanical Turk (142 females, 144 males, 2 others, 9 unknown; 46.86 ± 0.89 years old) between May 3 and July 2, 2024, using the CloudResearch Approved Group Filter. They received a base payment of $15.00 for completing the patch-leaving task, with a potential bonus of up to $5.00 contingent on performance on two randomly selected planets. Of these participants, 263 also filled out the Questionnaire of Unpredictability in Childhood (QUIC), and 147 filled out the Post-Traumatic Stress Disorder Checklist for DSM-5 (PCL-5), for an additional $5.00 compensation. The recruitment was limited to participants who (1) had a minimum 95% approval rate, (2) had completed at least 50 approved tasks, and (3) were located in the United States. Inclusion criteria required that participants (1) passed the instruction quiz with fewer than three attempts, (2) passed both attention checks, (3) had a mean residence time within two standard deviations of the group average, and (4) spent on average at least two digs per planet to ensure sufficient exposure to patch types.

### Data Analysis

Model fitting and parameter recovery analyses were performed in Julia 1.11.7. All subsequent data processing, statistical analyses, and visualization were performed in Python 3.12.4. To avoid assuming linear relationships or normally distributed variables, correlation analyses were conducted using Spearman’s rank correlation coefficient unless otherwise specified.

### Experiment Setup

Participants completed an online serial stay–switch foraging task adapted from prior human foraging tasks (e.g. (Constantino and Daw, 2015)). The primary way that the task differs from previous work is that it elicits behavior in the presence of multiple patch types (Harhen and Bornstein, 2023; Harhen et al., 2026; Chen et al., 2026). Participants were instructed to maximize total reward by traveling between planets and mining “space gems” across five blocks of fixed duration (6 minutes each), separated by self-paced breaks of up to one minute.

Upon arrival at a new planet (Fig. 1A), participants performed an initial dig and received a gem reward sampled from a Gaussian distribution (mean = 100, SD = 5). After each dig, they decided whether to stay on the current planet or leave to travel to a new one. Staying allowed immediate continuation of access to the current planet’s (depleting) resources, whereas leaving incurred a longer travel delay but granted access to a replenished gem mine. Participants had up to 2 seconds to respond; missed responses resulted in a brief timeout.

**Figure 1:**
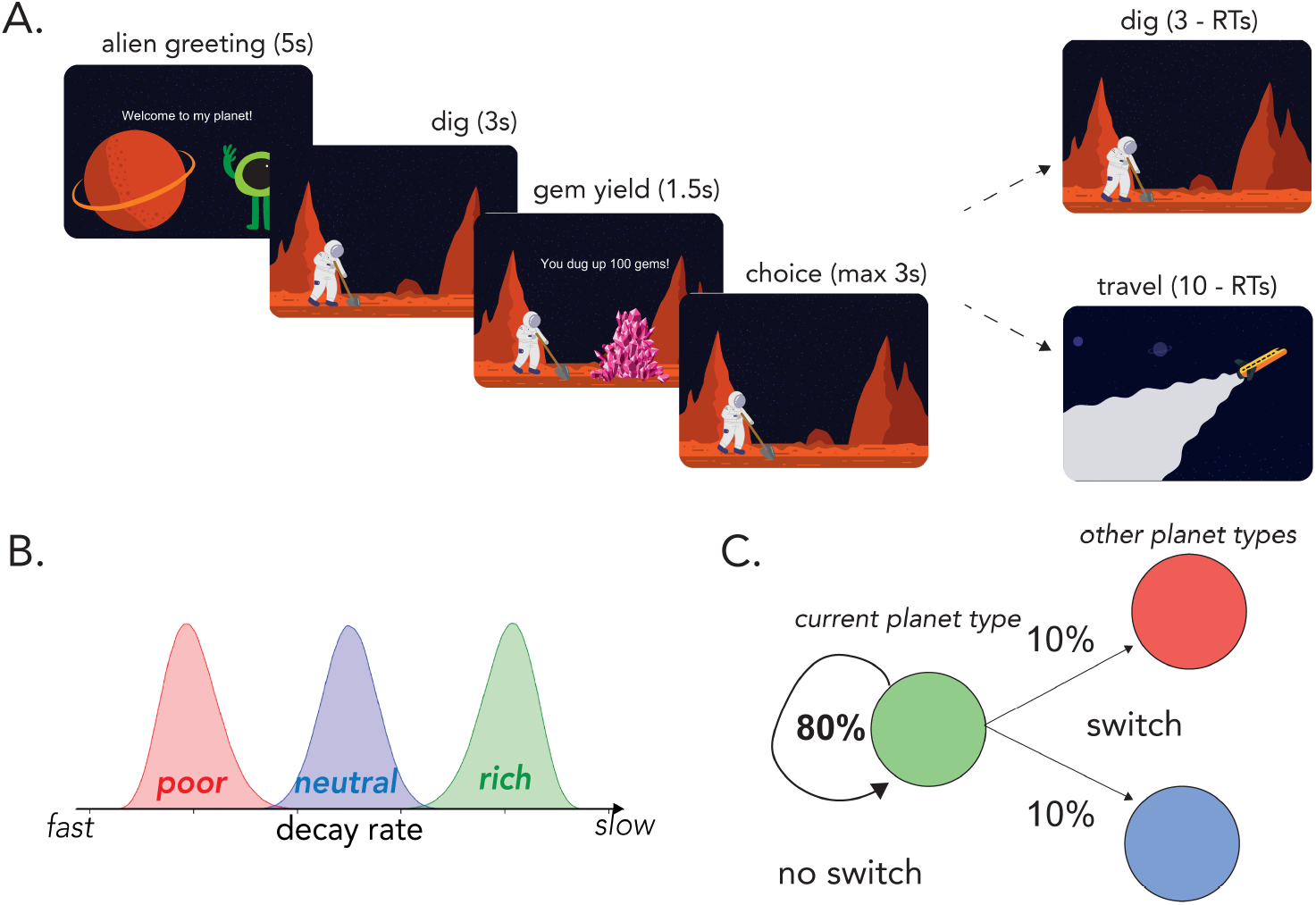
Planet Foraging Task: (A) Serial stay-switch task. Participants traveled to different planets and mined space gems across five 6-min blocks. On each trial, they had to decide between staying to dig from a depleting gem mine or incurring a time cost to travel to a new planet. (B) Environment structure. Planets varied in their richness or, more specifically, the rate at which they exponentially decayed with each dig. There were three planet types: poor, neutral, and rich—each with its own characteristic distribution over decay rates. (C) Environment dynamics. Planets of a similar type clustered together. A new planet had an 0.8 probability of being the same type as the prior planet (“no switch”). However, there was a 0.2 probability of transitioning or “switching” to a planet of a different type–equally distributed among each different planet type. (Figure adapted with permission from Harhen & Bornstein (2023).)

**Figure 2:**
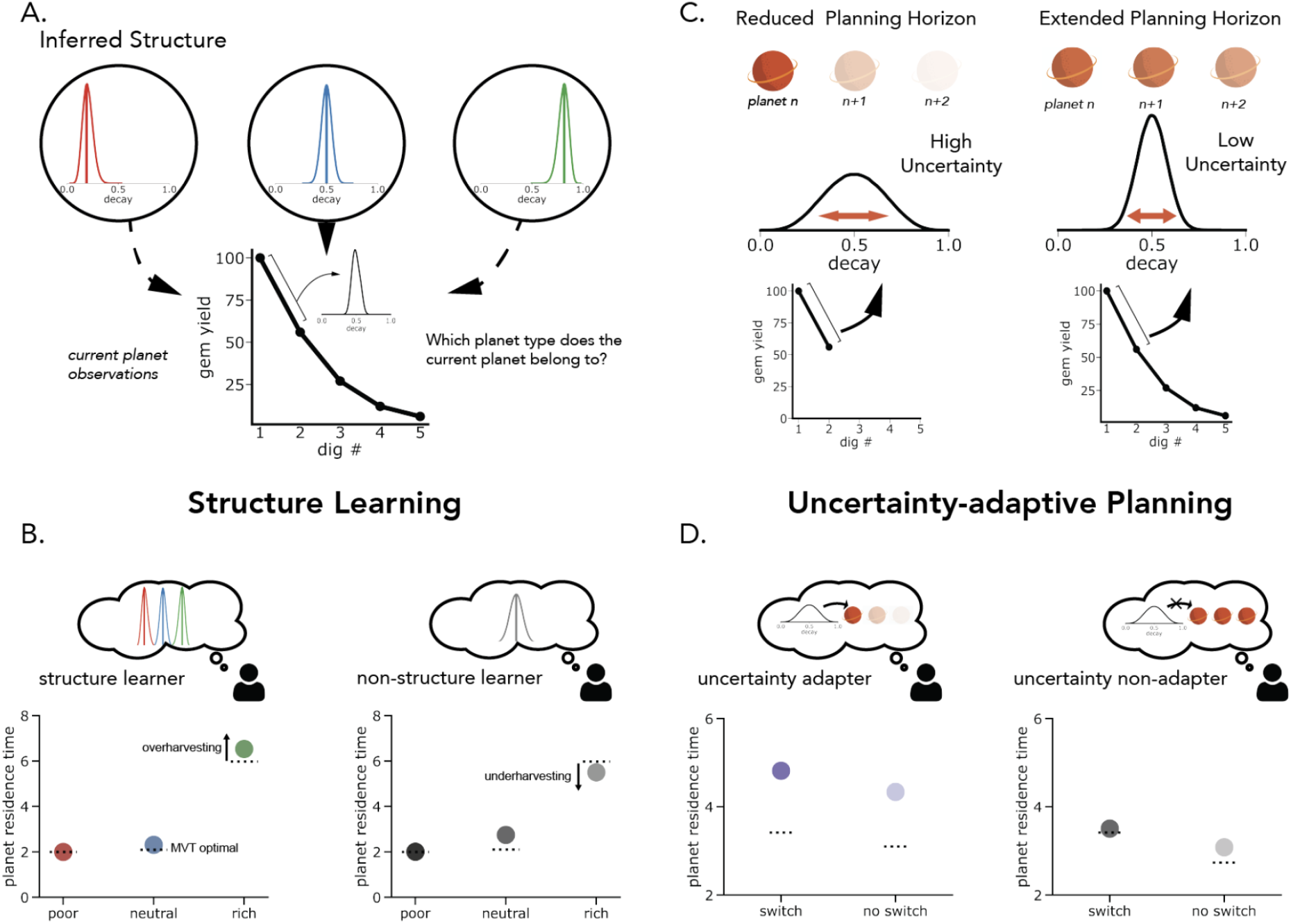
Computational framework for structure learning and uncertainty-dependent planning. (A) The forager performs two concurrent inferences: estimating the latent type of the current planet from experienced rewards and inferring the overall number of distinct planet types in the environment. We formalize this procedure as a Chinese Restaurant Process. (B) Model predictions from structure learning. The inferred complexity of the environment is controlled by the parameter *α*, which governs how readily new planet types are introduced. Different assumptions about environmental structure lead to systematic differences in harvesting behavior. Points denote simulated planet residence times (PRTs), and dashed lines indicate the Marginal Value Theorem (MVT) benchmark. (C) Uncertainty-dependent planning. The model assumes that the planning horizon is dynamically adjusted based on uncertainty about the current planet’s identity, with higher uncertainty promoting shorter planning horizons and lower uncertainty supporting longer-term planning. (D) Behavioral consequences of uncertainty modulation. Agents that adjust planning according to uncertainty exhibit increased overharvesting following infrequent transitions between planet types, whereas agents with fixed planning horizons show more stable residence times across environments. (Figure adapted with permission from Harhen et al., (2026).)

In order to isolate the effect of structure alone, temporal costs were controlled so that reaction time did not directly affect reward rate. Stay decisions were followed by a 1.5-second delay before reward feedback, with total trial duration held constant. Leave decisions incurred a 10-second travel delay, followed by arrival on a new planet before mining resumed.

Planets differed in richness, defined by the rate at which rewards decayed exponentially with successive digs. Three planet types were used—poor, neutral, and rich—with decay rates sampled from beta distributions producing fast, intermediate, and slow depletion, respectively (Fig. 1B).

Environmental structure extended across planets. When traveling to a new planet, there was an 80% probability that it would be of the same type as the previous planet and a 20% probability of switching to one of the other two types (Fig. 1C). This transition structure was not disclosed to participants.

### Optimal Planet Residence Time: Marginal Value Theorem

We compared participants’ planet residence time (PRT; the number of digs performed on the planet) to the optimal residence time prescribed by the Marginal Value Theorem (MVT). The MVT agent assumes perfect knowledge of decay rates and reward history.

The value of staying is:

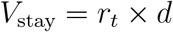

where *r*_*t*_ is the previous reward and *d* is the decay rate.

The value of leaving is:

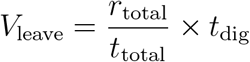

The agent chooses the higher of *V*_stay_ and *V*_leave_.

### Questionnaire of Unpredictability in Childhood (QUIC)

Participants completed the 38-item QUIC (Glynn et al., 2019), where each question assesses perceived unpredictability before ages 12 or 18. Responses were recorded as “yes,” “no,” or “prefer not to say.” The total score ranges from 0–38, with higher scores indicating greater early-life unpredictability.

To better characterize the latent structure of early life unpredictability in our sample, we conducted factor analysis on the 38 QUIC items, using a separate 732 individuals’ QUIC responses (278 females, 453 males, 1 other, 36.53 ± 10.99 years old, range 18-70) also collected on Amazon Mechanical Turk from May 2024 to July 2024 using the CloudResearch Approved Group filter. All items (38 in total) were included in the analysis. Maximum likelihood estimation was used for factor extraction, and an oblique rotation (Promax) was selected. Factors were thresholded by eigenvalue > 1. The EFA revealed that there are 6 latent factors. Full factor loadings and item assignments are provided in Supplementary Figure S3 and Table T1.

Items loading onto each factor were grouped accordingly, and factor scores were computed at the individual level by summing responses within each factor. These factor scores were standardized prior to analysis. The six factors were used in subsequent behavioral and computational analyses to examine whether specific dimensions of early-life unpredictability differentially predicted foraging behavior and model-derived parameters.

In addition to the factor-level analyses, we computed the total QUIC score as a global measure of early-life unpredictability to assess broad associations with behavioral and computational outcomes.

### Post-Traumatic Stress Disorder Checklist (PCL-5)

PCL-5 is a 20-item self-report measure of PTSD symptoms that corresponds to the criteria of DSM-5 (Wortmann et al., 2016). Participants rated symptom severity from 0 (“Not at all”) to 4 (“Extremely”). Total scores range from 0–80. We performed a factor analysis using the same procedure as that for QUIC, under which we identified three factors. Demographic associations with QUIC, PCL, and fitted model parameters are reported in Supplementary Figure S4.

### Adaptive Discounting Model

To characterize how participants inferred environmental structure and adjusted their decisions under uncertainty, we fit a Bayesian structure-learning and uncertainty-adaptive planning model adapted from recent work Harhen and Bornstein (2023); Harhen et al. (2026); Chen et al. (2026).

#### Structure Learning

Unlike the Marginal Value Theorem (MVT), which assumes perfect knowledge of patch types and their depletion dynamics, the current model assumes that foragers must infer the latent structure of the environment. Specifically, participants do not know (i) how many planet types exist, (ii) which planets belong to which type, or (iii) the depletion distribution governing each type.

We model structure inference using a Chinese Restaurant Process, CRP (Antoniak, 1974), a nonparametric Bayesian prior that allows the number of latent planet types to grow with experience. Under the CRP prior the probability that a newly encountered planet belongs to an existing cluster *k* is proportional to the number of previously assigned planets in that cluster:

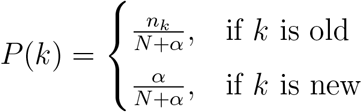

where *n*_*k*_ is the number of planets assigned to cluster *k, N* is the total number of planets encountered, and *α* governs the complexity of inferred structure. Larger values of *α* favor the creation of additional clusters.

After observing a sequence of depletions *D* on the current planet, the posterior probability that the planet belongs to cluster *k* is:

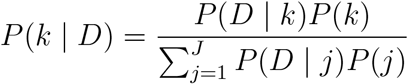

where *j* is the number of clusters formed thus far.

Because exact inference over cluster assignments is computationally intractable, we approximate the posterior using a particle filter (Djuric et al., 2003). Each particle represents a candidate clustering of previously encountered planets and is weighted according to its likelihood. Upon leaving a planet, particles are resampled in proportion to their weights, favoring representations that better explain observed reward dynamics. Each particle maintains a hypothetical clustering of planets, weighted by the likelihood of the data given that clustering. All simulations and fitting were done using a single particle, equivalent to Anderson’s local MAP algorithm (Anderson, 1991).

Each cluster maintains a Gaussian distribution over decay rates:

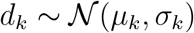

Cluster parameters are updated analytically using conjugate Normal–Gamma updates after each observed depletion.

To predict the next depletion, we use a Monte Carlo sampling procedure. For each simulation step: (1) a particle is sampled proportional to its weight, (2) a cluster is sampled from that particle’s posterior, and (3) a decay rate is drawn from the sampled cluster’s distribution. This process is repeated multiple times, and the sampled decay rates are averaged to generate the predicted depletion used for value computation.

#### Uncertainty-Adaptive Planning

Because structure inference is inherently uncertain, foragers must decide how far into the future to plan under epistemic uncertainty. Following prior work (Jiang et al., 2015; Harhen and Bornstein, 2023; Harhen et al., 2026; Chen et al., 2026), we model planning horizon adjustments via adaptive discounting of future rewards.

Representational uncertainty is quantified as the Shannon entropy of the multinomial distribution over planet types:

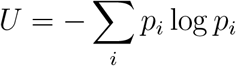

where *p*_*i*_ denotes the posterior probability of cluster *i*.

The effective discount factor governing planning depth is defined as a logistic function of baseline discounting and uncertainty sensitivity:

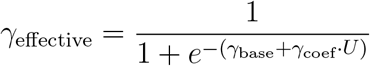

Here, *γ*_base_ captures baseline planning horizon, while *γ*_coef_ quantifies the extent to which structural uncertainty modulates discounting. Importantly, uncertainty in our task is not correlated with trial number (Supplementary Figure S5), ensuring that uncertainty-dependent effects cannot be attributed to time-on-task or learning progression.

#### Action Selection

Given predicted values for staying and leaving, decisions are generated via a softmax choice rule with lapse:

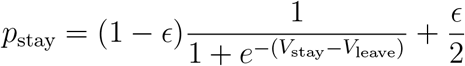

where *ϵ* is a lapse parameter capturing stimulus-independent random responses. The lapse term ensures that a small proportion of choices are made independently of the model’s value estimates, accounting for occasional inattentive or accidental responses.

This model dissociates two components of adaptive behavior: (1) the uncertainty of inferred environmental structure and (2) the extent to which uncertainty in that structure shapes planning depth. Together, these parameters allow us to test whether life experiences are associated with differences in structural representation, uncertainty-adaptive planning, or both.

### Parameter Recovery Analysis

To assess the identifiability of model parameters and the reliability of individual-differences inference, we conducted a parameter recovery analysis using simulated datasets. Parameter recovery evaluates whether the model-fitting procedure can recover known generative parameters and, critically for the present study, whether it preserves between-subject variability in those parameters.

#### Simulation Procedure

We simulated behavior for 200 agents performing the full task using the identical trial structure as human participants. For each simulated agent, parameters were independently drawn from broad uniform distributions spanning plausible ranges:

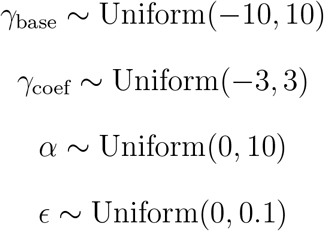

The simulation agents generated stay–leave decisions using the full adaptive discounting model, including CRP-based structure learning, particle filtering, uncertainty-dependent discounting, and softmax action selection with lapse.

#### Model Refitting

Each simulated dataset was refit using the same maximum likelihood estimation procedure applied to human participants. The fitting procedure estimated the three theoretically central parameters of the adaptive discounting model: the structure complexity parameter (*α*), baseline discounting (*γ*_base_), and uncertainty-sensitive discounting (*γ*_coef_). The lapse parameter (*ϵ*) was not estimated during fitting and was instead used only as simulation noise during data generation.

Because the primary goal of the present study is to examine individual differences (e.g. associations between model parameters and early-life unpredictability), our recovery evaluation focused on whether the fitting procedure preserves variability across agents. Recovery quality was quantified using Spearman correlation between true and fitted parameters across simulated agents.

#### Recovery Results

Baseline discounting (*γ*_base_) demonstrated robust recovery in rank space (Spearman *ρ* = 0.65; Supplementary Fig. S1B). Similarly, uncertainty sensitivity (*γ*_coef_) showed strong recovery (Spearman *ρ* = 0.71; Supplementary Fig. S1C), indicating that the model reliably preserves individual variability in baseline planning horizon and uncertainty-dependent discounting.

For the structure learning parameter *α*, we examined recovery after discretizing parameter values according to the modal number of inferred latent clusters produced by simulation of the CRP process. This is because the behavioral influence of *α* is mediated through the number of inferred latent clusters, which changes discretely rather than continuously as *α* varies. Consequently, small changes in *α* do not necessarily produce proportional changes in observed behavior, reducing the identifiability of exact continuous parameter values. (See Supplementary Fig. S1A for the comparatively poor recovery of *α* in continuous space.) Under this discretized representation, recovery performance was substantially improved (hit rate = 0.69; Supplementary Fig. S1D), indicating that although exact continuous values of *α* are difficult to identify, the model reliably captures coarse differences in inferred environmental structure complexity.

## Results

### Environmental richness and transition uncertainty shape harvesting behavior

First, we examined whether participants’ stay-leave behavior was sensitive to the reward structure of the task. We found that planet residence time (PRT) scaled with environmental richness: participants stayed longest in rich patches, intermediately in neutral patches, and shortest in poor patches (Fig. 3A). Pairwise comparisons confirmed robust richness-dependent adjustments in PRT (rich vs. neutral: *t*(296) = 30.54, *p* < .001; neutral vs. poor: *t*(296) = 19.18, *p* < .001). These effects indicate that participants tracked local reward statistics and adapted their stay/leave behavior accordingly.

**Figure 3:**
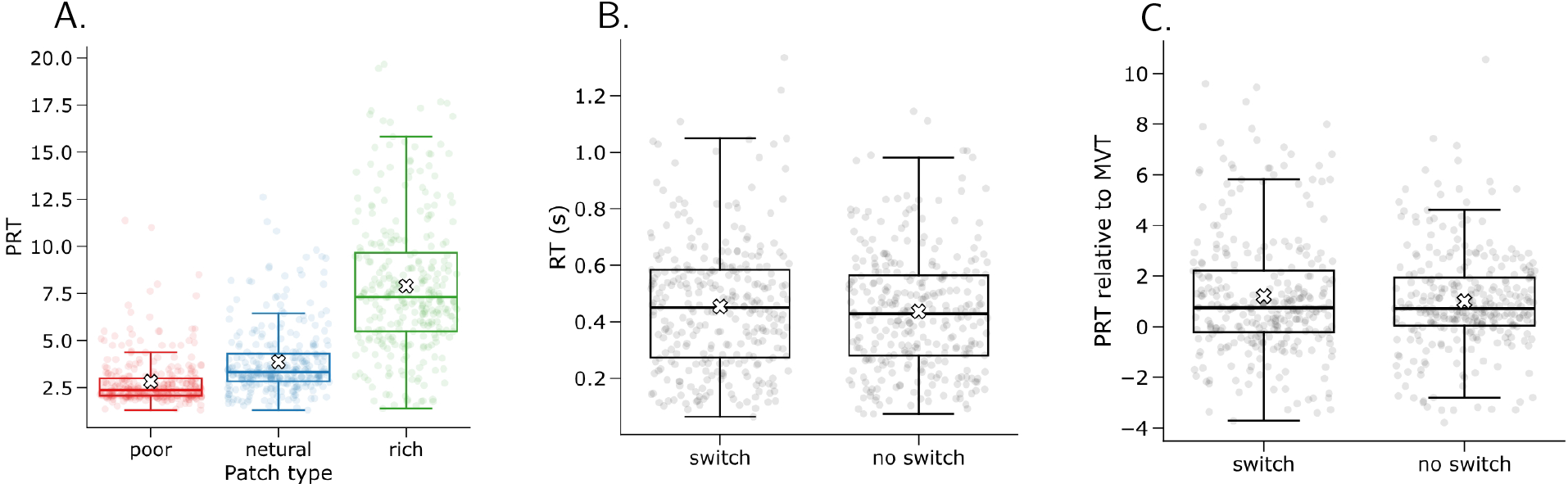
Environmental richness and transition uncertainty shape foraging behavior. (A) Mean planet residence time (PRT) differed systematically across patch types, with longer stays in richer environments. (B) Reaction times (RTs) were longer following rare patch-type switches than following common no-switch transitions. (C) Participants overharvested more following a switch, as indexed by larger PRT relative to the MVT optimum.

Next, we examined whether PRT performance approached optimality suggested by Marginal Value Theorem (MVT) with experience. Indeed, across the task, participants’ PRT moved closer to the MVT optimum as the absolute deviation between observed PRT and MVT decreased from the first to the final block (*t*(296) = −3.73, *p* < .001). This pattern suggests progressive rationalization of learning for environment’s reward structure.

A central prediction of our adaptive discounting model is that uncertainty should bias participants toward staying longer in the current patch by reducing the subjective value of leaving. To test this prediction behaviorally, we compared trials following rare patch-type switches to trials following common no-switch transitions. As expected, participants took longer to respond after a switch, consistent with increased decision uncertainty (Fig. 3B; *t*(296) = 3.40, *p* = .002). They also overharvested more following a switch, as indexed by higher PRT relative to the MVT optimum (Fig. 3C; *t*(296) = 3.73, *p* < .001). Together, these findings provide preliminary model-consistent behavioral evidence that uncertainty promotes longer staying.

We next asked whether individual differences in model parameters captured meaningful variation in foraging behavior and early-life unpredictability.

We first examined the relationship between early-life unpredictability (ELU) and uncertainty-derived parameters. Among the six ELU factors, only ELU Factor 1 showed a reliable association with the uncertainty-sensitive parameter r *γ*_coef_ (Fig. 4A; *r* = 0.18, *p* = .0024). This relationship remained significant after correction for multiple comparisons across all six ELU factors (Bonferroni-corrected *p* = .014), indicating that individuals reporting greater early-life unpredictability exhibit stronger trial-by-trial modulation of value by uncertainty.

**Figure 4:**
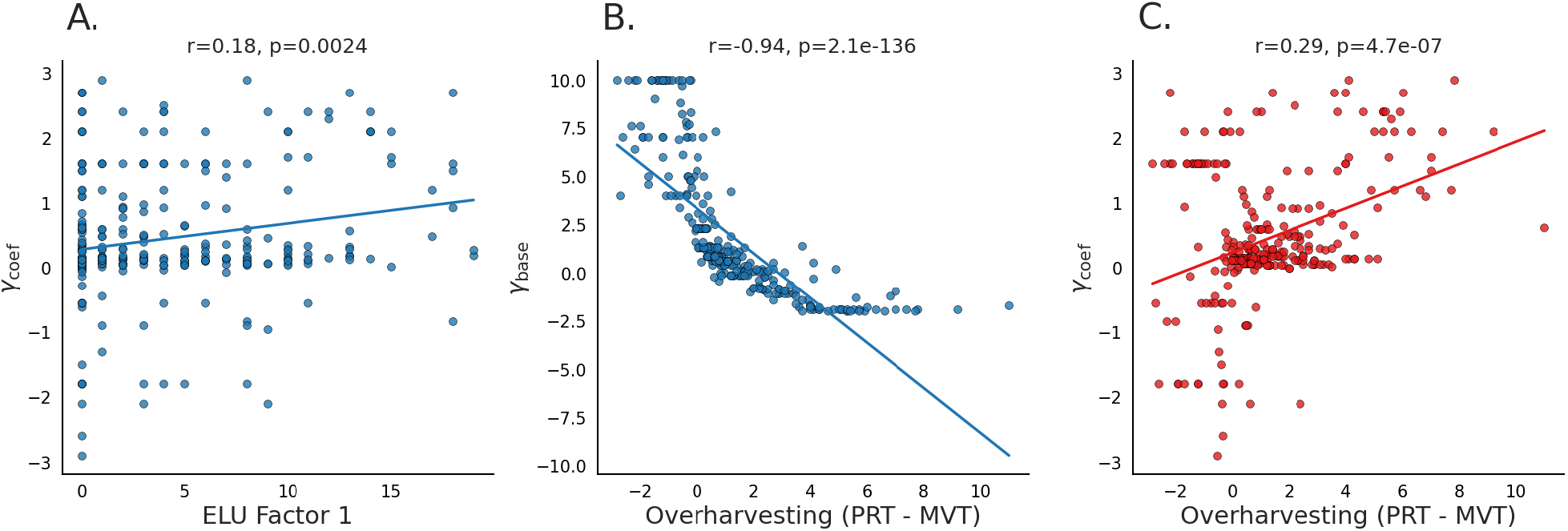
Behavioral deviations from optimal foraging relate to model-derived discounting parameters. (A) ELU Factor 1 was positively associated with *γ*_coef_, indicating stronger uncertainty-dependent modulation of leave valuation. (B) Greater overharvesting was strongly associated with lower *γ*_base_, indicating individual differences in discounting future rewards, irrespective of local uncertainty, drive overharvesting behavior. (C) Greater overharvesting was also associated with the simultaneously-fit higher *γ*_coef_ indicating stronger uncertainty-dependent modulation of leaving value, supporting the idea that individual differences in overharvesting were further modulated by local uncertainty.

We first examined how these computational parameters were related to deviations from optimal foraging i.e. overharvesting. Overharvesting, defined as actual PRT relative to the PRT suggested by Marginal Value Theorem (MVT), was strongly associated with the baseline discounting parameter *γ*_base_ (Fig. 4B; *r* = −0.94, *p* = 2.1 × 10^−136^), confirming that individuals who discounted future values were more likely to stay in the current patch. Importantly, additional variation in overharvesting was also associated accounted for by the uncertainty-sensitive component of the discount factor, *γ*_coef_ (Fig. 4C; *r* = 0.29, *p* = 4.7 × 10^−7^), linking a computational signature of subjective uncertainty weighting to a directly observable behavioral consequence—namely, staying longer than optimal. We observed a wide range of individual differences: distributions of model-derived parameters and behavioral measures are shown in Supplementary Figure S2.

We then asked whether ELU directly explained some of this variation in behavioral deviations from optimal foraging.

Specifically, given the robust associations between ELU and *γ*_coef_ (Fig. 4A) and between *γ*_coef_ and overharvesting (Fig. 4C), we tested whether ELU influenced behavior indirectly. Single-mediator analyses revealed a consistent indirect effect of ELU on overharvesting via *γ*_coef_ across environments. For poor environments, ELU significantly predicted overharvesting (total effect: *c* = 0.0520, *p* = .0011), and this effect was partially mediated by *γ*_coef_ (indirect effect: *ab* = 0.0216, 95% CI [0.0084, 0.0362]), with the direct effect remaining significant (*c*^*′*^ = 0.0304, *p* = .040). For neutral environments, the total effect was significant (*c* = 0.0492, *p* = .025), but the direct effect was not (*c*^*′*^ = 0.0213, *p* = .305), indicating full mediation (indirect effect: *ab* = 0.0279, 95% CI [0.0100, 0.0491]). For rich environments, neither the total nor direct effects were significant (*c* = 0.0493, *p* = .230; *c*^*′*^ = 0.0051, *p* = .898), but the indirect effect remained robust (indirect effect: *ab* = 0.0442, 95% CI [0.0155, 0.0792]), consistent with an indirect pattern only.

To see whether these effects generalized across environments, we next performed a single mediation analysis across all planet types at the subject level. Across all environments, ELU showed significant total effect on predicting overharvesting (total effect: *c* = 0.0502, *p* = .0022). Further, the indirect effect through *γ*_coef_ was significant (indirect effect: *ab* = 0.0311, 95% CI [0.0127, 0.0591]), while the direct effect was not significant after accounting for *γ*_coef_ (*c*^*′*^ = 0.0155, *p* = .528). Parallel mediation analyses that included both *γ*_coef_ and *γ*_base_ further showed that only the indirect pathway through *γ*_coef_ was significant (indirect effect via *γ*_coef_: 0.0246, 95% CI [0.0094, 0.0390]), whereas the indirect pathway through *γ*_base_ was not significant (indirect effect via *γ*_base_: 0.0111, 95% CI [−0.0204, 0.0463]).

Together, these results indicate that early-life unpredictability influences behavior through a latent computational pathway, consistent with theoretical proposals (Harhen and Bornstein, 2024). Specifically, ELU is associated with increased uncertainty-sensitive discounting, which in turn predicts overharvesting relative to the MVT benchmark. Consistent with this interpretation, ELU showed only a weak direct association with aggregate overharvesting, but the indirect pathway through uncertainty-sensitive discounting remained significant. This result emphasizes the influence early-life unpredictability on specific computational components of planning and valuation, even when its direct relationship with observed behavior is modest.

### Early-life unpredictability, but not PTSD symptom severity, predicts uncertainty weighting

Next, we examined whether individual differences in early-life unpredictability (ELU) and Post-Traumatic Stress Disorder Checklist (PCL) were associated with the uncertainty-weighting parameter, *γ*_coef_.

As with ELU, *γ*_coef_ was positively correlated with the total PCL score (Fig. 5A; *r* = 0.26, *p* = 1.8 × 10^−3^), suggesting that individuals with greater trauma-related symptoms exhibited a stronger modulation dependent on online uncertainty. However, PCL total was also strongly correlated with overall ELU measured by QUIC total (Fig. 5B; *r* = 0.50, *p* = 1.1 × 10^−10^), indicating substantial shared variance.

**Figure 5:**
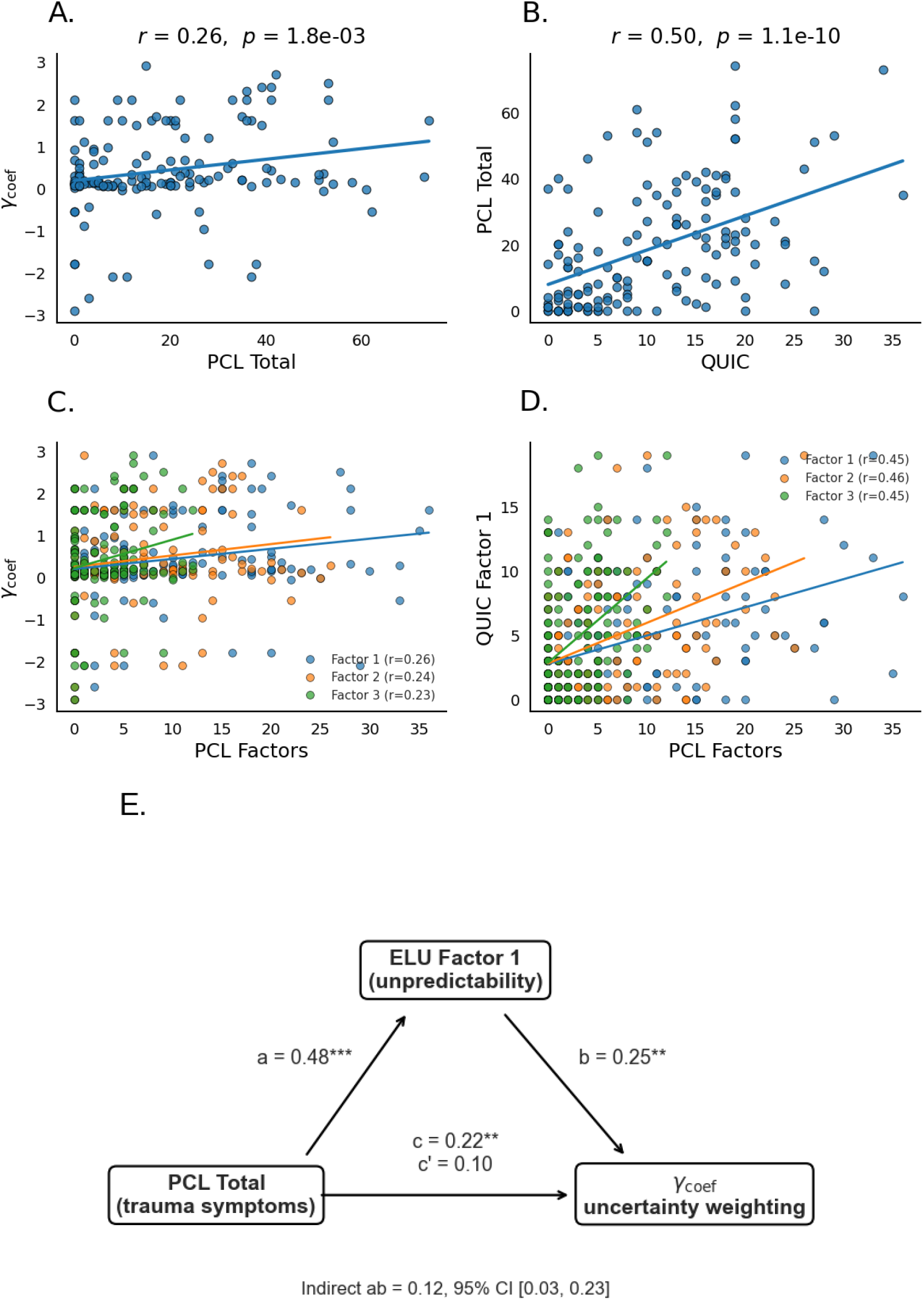
Dissociation between early-life unpredictability and PTSD symptoms in predicting uncertainty weighting. (A) *γ*_coef_ is positively correlated with PCL total. (B) PCL total is strongly correlated with QUIC total. (C) *γ*_coef_ shows weak positive correlations with PCL-derived factors. (D) PCL-derived factors are moderately correlated with ELU Factor 1. (E) Mediation analysis testing whether the association between PCL total and *γ*_coef_ is mediated by ELU Factor 1.

To further elucidate this relationship, we examined factor-level structure. *γ*_coef_ showed small positive correlations with each of the three PCL-derived factors (Fig. 5C; *r*s ≈ 0.23– 0.26). Importantly, ELU Factor 1 was positively correlated with all PCL factors (Fig. 5D): PCL Factor 1 (*r* = 0.45, *p* = 8.52 × 10^−8^), PCL Factor 2 (*r* = 0.46, *p* = 1.65 × 10^−8^), and PCL Factor 3 (*r* = 0.45, *p* = 8.05 × 10^−9^). These results indicate that greater early-life unpredictability was broadly associated with elevated trauma-related symptom dimensions.

To determine whether ELU or PTSD symptom severity uniquely explained uncertainty weighting, we next performed multiple regression analyses including both ELU and PCL measures as simultaneous predictors of *γ*_coef_. When ELU Factor 1 and total PCL score were entered together into the same model, the overall regression was significant (*R*^2^ = 0.10, *F* (2, 144) = 7.77, *p* = 6.24 × 10^−4^). However, only ELU Factor 1 uniquely predicted *γ*_coef_ (*β* = 0.25, *p* = .006), whereas PCL total no longer explained unique variance (*β* = 0.10, *p* = .262). Similarly, when ELU Factor 1 and all three PCL-derived factors were included simultaneously, the overall model remained significant (*R*^2^ = 0.10, *F* (4, 142) = 4.09, *p* = .0036), but again only ELU Factor 1 significantly predicted *γ*_coef_ (*β* = 0.25, *p* = .007), whereas none of the individual PCL factors were significant predictors (all *p*s > .38).

We further tested whether the apparent relationship between PTSD symptoms and uncertainty weighting was mediated by ELU (Fig. 5E). Mediation analyses showed that PCL total significantly predicted ELU Factor 1 (path *a*: *β* = 0.48, *p <* .001), and ELU Factor 1 significantly predicted *γ*_coef_ after controlling for PCL (path *b*: *β* = 0.25, *p* = .006). Although PCL total initially predicted *γ*_coef_ (total effect *c*: *β* = 0.22, *p* = .007), this direct relationship was no longer significant after accounting for ELU (direct effect *c*^*′*^: *β* = 0.10, *p* = .262). The indirect effect through ELU was significant (indirect effect *ab* = 0.12, 95% bootstrap CI [0.03, 0.23]), indicating that the association between PTSD symptom severity and uncertainty weighting was largely explained by variance shared with ELU.

In summary, the apparent relationship between PTSD symptom severity and uncertainty weighting is accounted for by variance shared with early-life unpredictability. In contrast, ELU uniquely predicts individual differences in adaptive uncertainty discounting, indicating that online uncertainty-sensitive discounting is more closely related to developmental unpredictability than to the severity of current trauma symptoms. This dissociation between early-life unpredictability and lifelong trauma experience supports the idea that the influence of unpredictability obtains during a sensitive period of plasticity during early development (Birnie and Baram, 2022; Harhen and Bornstein, 2024), unlike later traumas.

## Discussion

In our present study, we investigated how early-life unpredictability (ELU) shape sequential stay-leave decisions under uncertainty. Using a structured patch foraging task and a computational model of uncertainty-adaptive planning, we showed that ELU was associated with uncertainty sensitivity discounting. Individuals reporting higher ELU exhibited stronger uncertainty sensitivity for environmental structure that in terms modulates stay-leave behavior. Importantly, this uncertainty sensitivity predicts systematic overharvesting relative to the Marginal Value Theorem (MVT) benchmark, and mediation analyses showed that the relationship between ELU and overharvesting was carried indirectly through uncertainty sensitivity. Together, our findings suggest that early-life unpredictability shapes adult decision-making not by directly altering observable behavior, but by tuning how uncertainty influences planning and valuation.

Our findings build on recent theoretical work proposing that unpredictability during developmental-sensitive period selectively alters psychiatric symptoms and neural systems involved in representing uncertainty and future expectations (Baram et al., 2012; Birnie and Baram, 2022; Spadoni et al., 2022). We identify a computational pathway that links ELU to behavior in a naturalistic foraging task. In our model, uncertainty reduces the effective planning horizon by discounting the value of leaving a patch. Participants with greater ELU therefore behaved as if uncertain future reward were less valuable, biasing them toward overharvesting in current patch. With our structural learning modeling, we replicate and extend prior work by showing that human overharvesting can emerge from rational adaptation to latent environmental uncertainty rather than from simple failures of self-control or impulsivity (Constantino and Daw, 2015; Harhen and Bornstein, 2023; Harhen et al., 2026; Chen et al., 2026). Within this framework, overharvesting reflects a computationally meaningful consequence of uncertainty-sensitive planning. Our results suggest that ELU systematically shifts the degree to which uncertainty shapes future-oriented valuation.

A particularly interesting aspect of the present findings is the dissociation between early-life unpredictability and trauma-related symptom severity. Although PTSD symptoms initially showed positive associations with uncertainty weighting, these effects were accounted for by variance shared with ELU. Once ELU and PCL factors were modeled simultaneously, only ELU uniquely predicted uncertainty sensitivity. This dissociation suggests that developmental unpredictability may influence uncertainty-sensitive planning through mechanisms partially distinct from those associated with later traumatic experiences or current symptom burden. More broadly, the findings support emerging perspectives arguing that unpredictability during sensitive developmental periods calibrates core inferential and valuation systems that persist into adulthood (Birnie and Baram, 2022).

For future research directions, our work raises the possibility that ELU shapes neural systems involved in predictive representations and uncertainty estimation (Stachenfeld et al., 2017; Duvelle et al., 2023; Shenhav et al., 2014; Hayden et al., 2011). Sequential foraging requires maintaining beliefs about hidden environmental structures, estimating transition uncertainty, and planning for future actions.

Future work could extend this framework to environments in which participants voluntarily choose transitions among distinct patch types, where choices may be influenced by uncertainty sensitivity (Charnov, 1976). Further, combining computational modeling with neuroimaging could clarify how ELU shapes representations of environment states–are these states merged in individuals with higher subjective uncertainty, or does uncertainty modulation affect the value computations alone?

Our study is not without limitations. First, the present data cannot establish causal developmental effects of ELU in directly shaping uncertainty-sensitive planning. To measure that, longitudinal studies will be needed. Similarly, the current study relied on retrospective self-report measures of unpredictability and trauma exposure, which may be influenced by recall biases (Zhang et al., 2026).

In summary, we show that early-life unpredictability is associated with increased uncertainty-sensitive discounting during sequential foraging. This latent computational phenotype predicts systematic overharvesting relative to normative foraging benchmarks and mediates the relationship between developmental unpredictability and behavior. By dissociating the influence of early-life unpredictability from trauma symptom severity, the present work provides evidence that developmental unpredictability selectively shapes how uncertainty is incorporated into planning and valuation. These findings support the broader idea that early environments calibrate core computational mechanisms underlying adaptive behavior under uncertainty.

## Data Availability

All data and code can be found in the GitHub link: https://github.com/zyxl100/foraging_MQLP_2026

## Acknowledgments

AMB acknowledges funding from NIMH (R01MH061285, PI: SJ Pollak; P50MH096889, PI: TZB).

## Author Contributions

## References

Anderson, J. R. (1991). The adaptive nature of human categorization. Psychological review, 98(3):409.

Antoniak, C. E. (1974). Mixtures of dirichlet processes with applications to bayesian non-parametric problems. The annals of statistics, pages 1152–1174.

Baram, T. Z., Davis, E. P., Obenaus, A., Sandman, C. A., Small, S. L., Solodkin, A., and Stern, H. (2012). Fragmentation and unpredictability of early-life experience in mental disorders. American Journal of Psychiatry, 169(9):907–915.

Birnie, M. T. and Baram, T. Z. (2022). Principles of emotional brain circuit maturation. Science, 376(6597):1055–1056.

Blanchard, T. C. and Hayden, B. Y. (2015). Monkeys are more patient in a foraging task than in a standard intertemporal choice task. PloS one, 10(2):e0117057.

Charnov, E. L. (1976). Optimal foraging, the marginal value theorem. Theoretical population biology, 9(2):129–136.

Chen, X., Johnson, M., Ra, P., Maxwell, M., Harhen, N., Noh, S., Bennett, I., and Bornstein, A. (2026). Age-related differences in structure learning drive foraging behavior in a multimodal patch-leaving task. In Cognitive Computational Neuroscience.

Constantino, S. M. and Daw, N. D. (2015). Learning the opportunity cost of time in a patch-foraging task. Cognitive, Affective, & Behavioral Neuroscience, 15(4):837–853.

De Berker, A. O., Rutledge, R. B., Mathys, C., Marshall, L., Cross, G. F., Dolan, R. J., and Bestmann, S. (2016). Computations of uncertainty mediate acute stress responses in humans. Nature communications, 7(1):10996.

Decker, A. L., Leonard, J., Romeo, R., Itiat, J., Hubbard, N. A., Bauer, C. C., Grotzinger, H., Giebler, M. A., Torres, Y. C., Imhof, A., et al. (2025). Exploration is associated with socioeconomic disparities in learning and academic achievement in adolescence. Nature Communications, 16(1):6342.

Djuric, P. M., Kotecha, J. H., Zhang, J., Huang, Y., Ghirmai, T., Bugallo, M. F., and Miguez, J. (2003). Particle filtering. IEEE signal processing magazine, 20(5):19–38.

Duvelle, É., Grieves, R. M., and Van der Meer, M. A. (2023). Temporal context and latent state inference in the hippocampal splitter signal. Elife, 12:e82357.

Glynn, L. M., Stern, H. S., Howland, M. A., Risbrough, V. B., Baker, D. G., Nievergelt, C. M., Baram, T. Z., and Davis, E. P. (2019). Measuring novel antecedents of mental illness: the questionnaire of unpredictability in childhood. Neuropsychopharmacology, 44(5):876–882.

Harhen, N. C. and Bornstein, A. M. (2023). Overharvesting in human patch foraging reflects rational structure learning and adaptive planning. Proceedings of the National Academy of Sciences, 120(13):e2216524120.

Harhen, N. C. and Bornstein, A. M. (2024). Interval timing as a computational pathway from early life adversity to affective disorders. Topics in cognitive science, 16(1):92–112.

Harhen, N. C., Budiono, R., Hartley, C. A., and Bornstein, A. M. (2026). Structure inference in complex environments improves from childhood to adulthood. Developmental Science, 29(3):e70163.

Hayden, B. Y., Pearson, J. M., and Platt, M. L. (2011). Neuronal basis of sequential foraging decisions in a patchy environment. Nature neuroscience, 14(7):933–939.

Jiang, N., Kulesza, A., Singh, S., and Lewis, R. (2015). The dependence of effective planning horizon on model accuracy. In Proceedings of the 2015 international conference on autonomous agents and multiagent systems, pages 1181–1189.

Morgado, P., Sousa, N., and Cerqueira, J. J. (2015). The impact of stress in decision making in the context of uncertainty. Journal of Neuroscience Research, 93(6):839–847.

Rachum, A., Harten, L. M., Assa, R., Goldshtein, A., Chen, X., Gonceer, N., and Yovel, Y. (2025). Early experience affects foraging behavior of wild fruit bats more than their original behavioral predispositions. eLife, 14:RP103220.

Shenhav, A., Straccia, M. A., Cohen, J. D., and Botvinick, M. M. (2014). Anterior cingulate engagement in a foraging context reflects choice difficulty, not foraging value. Nature neuroscience, 17(9):1249–1254.

Spadoni, A. D., Vinograd, M., Cuccurazzu, B., Torres, K., Glynn, L. M., Davis, E. P., Baram, T. Z., Baker, D. G., Nievergelt, C. M., and Risbrough, V. B. (2022). Contribution of early-life unpredictability to neuropsychiatric symptom patterns in adulthood. Depression and anxiety, 39(10-11):706–717.

Stachenfeld, K. L., Botvinick, M. M., and Gershman, S. J. (2017). The hippocampus as a predictive map. Nature neuroscience, 20(11):1643–1653.

Verfaellie, M., Patt, V., Lafleche, G., Jones, D., and Vasterling, J. J. (2025). Choosing certainty over risk: Associations of ptsd symptom severity with memory sampling during experiential decision making. Journal of Anxiety Disorders, 110:102979.

Wortmann, J. H., Jordan, A. H., Weathers, F. W., Resick, P. A., Dondanville, K. A., Hall-Clark, B., Foa, E. B., Young-McCaughan, S., Yarvis, J. S., Hembree, E. A., et al. (2016). Psychometric analysis of the ptsd checklist-5 (pcl-5) among treatment-seeking military service members. Psychological assessment, 28(11):1392.

Zhang, Z., Zhou, C., Zhang, R., Tang, Y., Zhang, Y., Qin, P., Su, B., and Wang, Y. (2026). Depression shapes the recall of adverse childhood experiences: evidence from a three-wave longitudinal study of 6,260 chinese adolescents. Nature Mental Health, pages 1–12.

