## Supplementary material for "A specific computational role for early-life unpredictability, and not lifelong traumatic experience, in decision-making under uncertainty"

### Supplementary Materials

### S1. Model Parameter Recovery

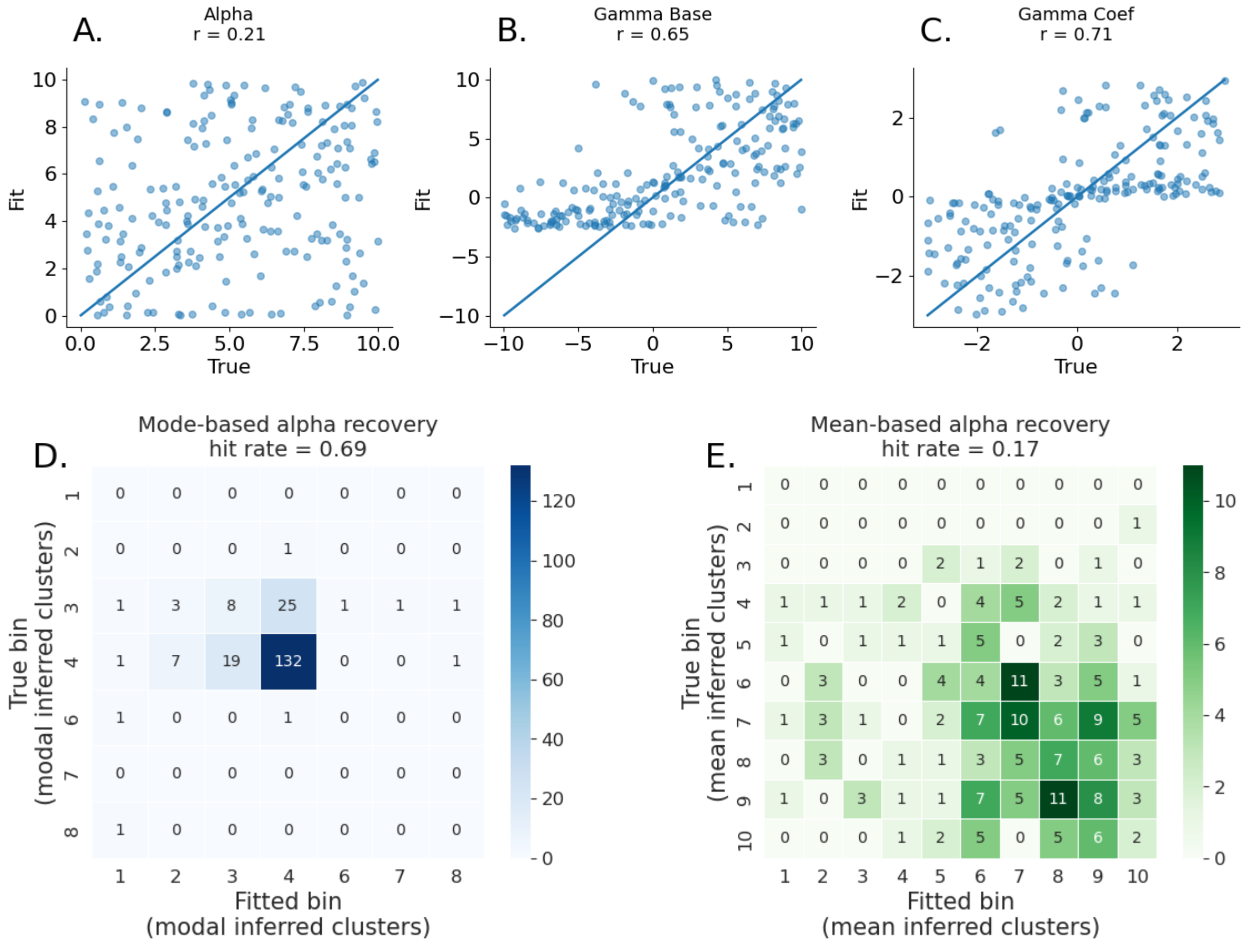


Figure S1: **Parameter recovery of the adaptive discounting model with 200 simulated subjects.** (A) Structure complexity parameter ($\alpha$) shows weak continuous recovery, reflecting the coarse and nonlinear relationship between $\alpha$ and inferred latent structure. (B) Baseline discounting ($\gamma_{\text{base}}$) is robustly recovered, indicating reliable estimation of individual differences in baseline planning horizon. (C) Uncertainty sensitivity ($\gamma_{\text{coef}}$) is strongly recovered, supporting its interpretability as a key individual-differences parameter. (D) Recovery of $\alpha$ in discretized modal-cluster space. True and fitted $\alpha$ values were mapped to the modal number of inferred latent clusters generated by CRP simulations, with $\alpha$ values discretized while all other model parameters were uniformly sampled. Recovery accuracy was quantified as the proportion of simulated subjects whose fitted and true modal cluster bins matched (hit rate = 0.69), indicating reliable recovery of coarse differences in inferred structural complexity. (E) Recovery of $\alpha$ in discretized mean-cluster space. True and fitted $\alpha$ values were mapped to bins that are defined as rounded mean number of inferred latent clusters. Recovery accuracy was weaker in this average representation (hit rate = 0.17).

#### S2. Fitted Parameter, QUIC, and PCL Dsitribution

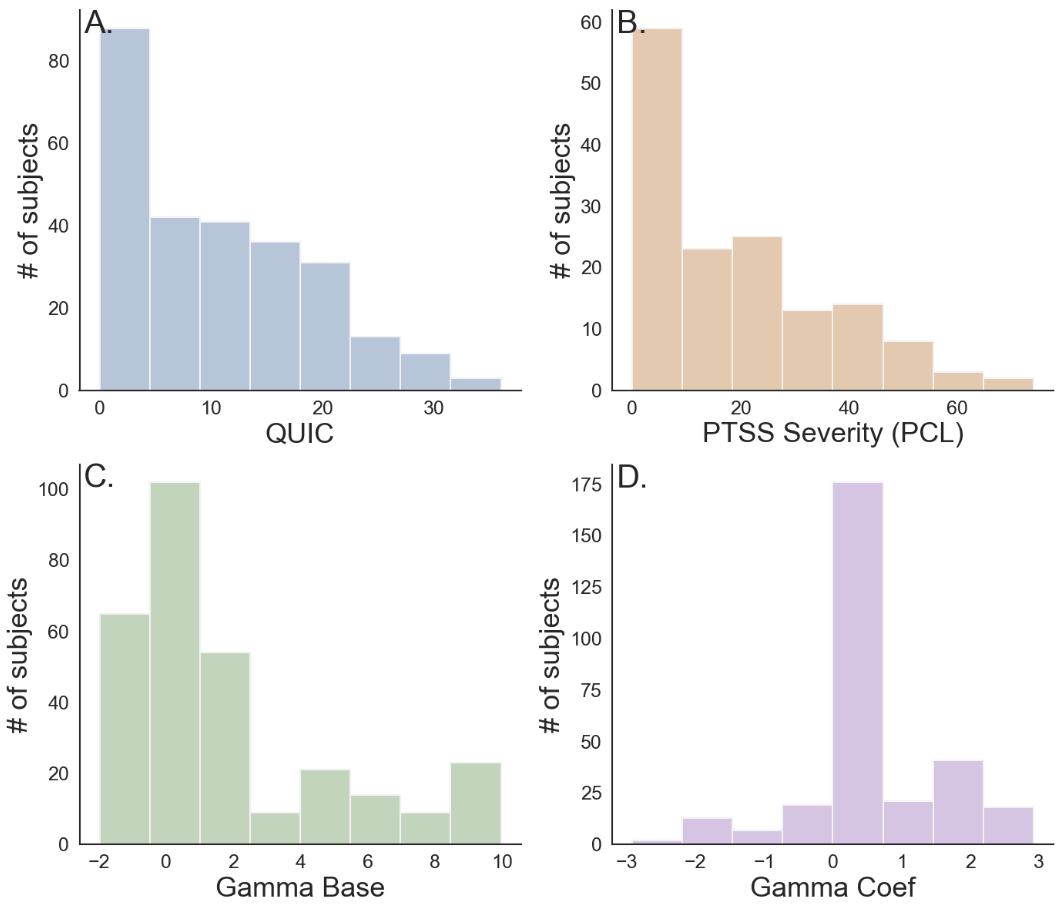


**Figure S2: Distribution of behavioral and model-derived variables.** (A) Distribution of total early-life unpredictability scores (QUIC), showing variability across participants. (B) Distribution of PTSD symptom severity (PCL total), showing a wide range of reported symptoms. (C) Distribution of baseline discounting parameter ($\gamma_{\text{base}}$), reflecting individual differences in baseline planning horizon. (D) Distribution of uncertainty sensitivity parameter ($\gamma_{\text{coef}}$), reflecting individual differences in trial-by-trial uncertainty-dependent modulation.

### S3. Factor Analysis Details

| | Factor 1 | Factor 2 | Factor 3 | Factor 4 | Factor 5 | Factor 6 | Uniqueness |
|---|---|---|---|---|---|---|---|
| Q12 | **0.841** | 0.007 | −0.019 | −0.078 | −0.013 | −0.140 | 0.395 |
| Q23 | **0.790** | 0.073 | −0.057 | 0.012 | −0.023 | 0.055 | 0.383 |
| Q13 | **0.750** | 0.020 | 0.011 | −0.060 | 0.003 | −0.013 | 0.470 |
| Q21 | **0.738** | 0.095 | 0.036 | 0.062 | −0.058 | −0.136 | 0.448 |
| Q2 | **0.685** | 0.047 | 0.014 | −0.102 | −0.034 | 0.020 | 0.563 |
| Q19 | **0.637** | −0.033 | 0.005 | 0.096 | −0.018 | −0.050 | 0.549 |
| Q20 | **0.624** | 0.025 | −0.113 | **0.379** | −0.035 | −0.127 | 0.417 |
| Q25 | **0.601** | 0.180 | 0.232 | −0.043 | −0.102 | −0.007 | 0.473 |
| Q24 | **0.589** | 0.039 | 0.106 | 0.079 | 0.060 | 0.038 | 0.503 |
| Q16 | **0.585** | −0.031 | 0.183 | −0.056 | −0.027 | 0.034 | 0.543 |
| Q22 | **0.582** | −0.029 | 0.056 | 0.091 | 0.055 | −0.018 | 0.576 |
| Q37 | **0.509** | −0.121 | −0.052 | −0.008 | 0.072 | **0.529** | 0.357 |
| Q38 | **0.505** | −0.102 | 0.036 | −0.011 | 0.034 | **0.419** | 0.431 |
| Q30 | **0.490** | 0.008 | 0.102 | 0.180 | −0.008 | 0.028 | 0.574 |
| Q26 | **0.488** | −0.075 | −0.141 | **0.559** | 0.023 | −0.068 | 0.372 |
| Q18 | **0.483** | −0.077 | 0.083 | 0.227 | 0.109 | −0.060 | 0.576 |
| Q27 | **0.335** | −0.033 | −0.097 | **0.593** | 0.001 | −0.009 | 0.462 |
| Q11 | **0.331** | −0.113 | **0.461** | −0.117 | 0.103 | 0.008 | 0.586 |
| Q28 | **−0.318** | **0.358** | 0.215 | 0.238 | 0.023 | −0.010 | 0.658 |
| Q34 | 0.213 | −0.022 | **0.731** | −0.124 | 0.048 | −0.114 | 0.422 |
| Q8 | 0.194 | **0.463** | −0.075 | −0.025 | −0.029 | 0.061 | 0.773 |
| Q36 | −0.185 | 0.173 | −0.031 | 0.007 | 0.011 | **0.689** | 0.453 |
| Q1 | 0.182 | **0.427** | −0.108 | −0.053 | 0.017 | 0.010 | 0.820 |
| Q35 | 0.173 | 0.042 | **0.641** | −0.044 | −0.120 | −0.017 | 0.503 |
| Q5 | 0.170 | **0.695** | −0.120 | 0.012 | −0.049 | 0.007 | 0.566 |
| Q3 | −0.142 | **0.584** | 0.013 | 0.001 | −0.081 | 0.032 | 0.646 |
| Q15 | −0.132 | **0.538** | 0.152 | 0.028 | 0.048 | −0.060 | 0.631 |
| Q33 | 0.107 | −0.019 | **0.706** | 0.049 | −0.032 | −0.026 | 0.412 |
| Q7 | 0.100 | **0.509** | 0.002 | −0.026 | 0.099 | −0.052 | 0.711 |
| Q9 | −0.090 | **0.588** | 0.082 | −0.015 | 0.088 | −0.031 | 0.580 |
| Q31 | 0.090 | 0.084 | **0.314** | 0.181 | −0.079 | 0.243 | 0.597 |
| Q6 | −0.056 | **0.612** | 0.002 | −0.039 | −0.037 | −0.029 | 0.649 |
| Q14 | −0.032 | **0.580** | 0.117 | −0.077 | 0.125 | 0.015 | 0.542 |
| Q4 | −0.029 | **0.523** | −0.047 | 0.058 | −0.035 | 0.142 | 0.668 |
| Q29 | 0.029 | −0.013 | 0.135 | **0.422** | 0.008 | 0.050 | 0.721 |
| Q17 | −0.022 | 0.187 | −0.048 | 0.021 | **0.755** | 0.019 | 0.282 |
| Q32 | −0.003 | −0.003 | **0.692** | 0.108 | −0.009 | 0.027 | 0.429 |
| Q10 | 0.003 | **0.388** | −0.097 | 0.015 | **0.634** | 0.057 | 0.242 |

**Figure S3: Factor loadings of QUIC items.** Factor loadings from exploratory factor analysis of the 38 QUIC items. Each row corresponds to an individual item, and columns indicate loadings on each latent factor. Values reflect the strength of association between each item and the corresponding factor, with higher absolute values indicating stronger contributions. This structure was used to group items and compute factor-level scores for subsequent analyses.

Table S1: Factor structure and corresponding QUIC items.

| **Factor** | **Subscale** | **Question** |
|---|---|---|
| 1 | Parental predictability | 2: My parents were often late to pick me up (e.g., from school, aftercare, or sports). |
| 1 | Parental predictability | 11: At least one of my parents had punishments that were unpredictable. |
| 1 | Parental predictability | 12: I often wondered whether or not one of my parents would come home at the end of the day. |
| 1 | Physical environment | 13: There were often people coming and going in my house that I did not expect to be there. |
| 1 | Parental predictability | 16: At least one of my parents would plan something for the family, but then not follow through with the plan. |
| 1 | Parental environment | 18: There was a long period of time when I did not see one of my parents (e.g., military deployment, jail time, custody arrangements). |
| 1 | Parental environment | 19: I experienced changes in my custody arrangement. |
| 1 | Physical environment | 20: I moved frequently. |
| 1 | Parental environment | 21: At least one of my parents changed jobs frequently. |
| 1 | Parental environment | 22: There were times when one of my parents was unemployed and could not find a job even though he/she wanted one. |
| 1 | Safety and security | 23: There was a period of time when I often worried that I was not going to have enough food to eat. |

| Factor | Subscale | Question |
|---|---|---|
| 1 | Safety and security | 24: There was a period of time when I often worried that my family would not have enough money to pay for necessities like clothing or bills. |
| 1 | Safety and security | 25: There was a period of time when I did not feel safe in my home. |
| 1 | Physical environment | 26: I changed schools frequently. |
| 1 | Physical environment | 27: I changed schools mid-year. |
| 1 | Parental environment | 28: My parents had a stable relationship with each other. |
| 1 | Parental environment | 30: At least one of my parents had many romantic partners. |
| 1 | Physical environment | 37: I lived in a cluttered house (e.g., piles of stuff everywhere). |
| 1 | Physical environment | 38: In my house, things I needed were often misplaced so that I could not find them. |
| 2 | Parental monitoring and involvement | 1: I had a set morning routine on school days (i.e., I usually did the same thing each day to get ready). |
| 2 | Parental monitoring and involvement | 3: My parents kept track of what I ate (e.g., made sure that I did not skip meals or tried to make sure I ate healthy food). |
| 2 | Parental monitoring and involvement | 4: My family ate a meal together most days. |
| 2 | Parental monitoring and involvement | 5: My parents tried to make sure I got a good night's sleep (e.g., I had a regular bedtime, my parents checked to make sure I went to sleep). |
| 2 | Parental monitoring and involvement | 6: I had a bedtime routine (e.g., my parents tucked me in, my parents read me a book, I took a bath). |
| 2 | Parental monitoring and involvement | 7: In my afterschool or free-time hours, at least one of my parents knew what I was doing. |
| 2 | Parental predictability | 8: I usually knew when my parents were going to be home. |

| Factor | Subscale | Question |
|---|---|---|
| 2 | Parental monitoring and involvement | 9: At least one of my parents regularly checked that I did my homework. |
| 2 | Parental monitoring and involvement | 10: At least one of my parents regularly kept track of my school progress. |
| 2 | Parental monitoring and involvement | 14: At least one parent made time each day to see how I was doing. |
| 2 | Parental predictability | 15: My family planned activities to do together. |
| 2 | Parental environment | 28: My parents had a stable relationship with each other. |
| 3 | Parental predictability | 11: At least one of my parents had punishments that were unpredictable. |
| 3 | Parental predictability | 31: At least one of my parents was disorganized. |
| 3 | Parental predictability | 32: At least one of my parents was unpredictable. |
| 3 | Parental predictability | 33: For at least one of my parents, when they were upset I did not know how they would act. |
| 3 | Parental predictability | 34: One of my parents could go from calm to furious in an instant. |
| 3 | Parental predictability | 35: One of my parents could go from calm to stressed or nervous in an instant. |
| 4 | Physical environment | 20: I moved frequently. |
| 4 | Physical environment | 26: I changed schools frequently. |
| 4 | Physical environment | 27: I changed schools mid-year. |
| 4 | Parental environment | 29: My parents got divorced. |
| 5 | Parental monitoring and involvement | 10: At least one of my parents regularly kept track of my school progress. |

| Factor | Subscale | Question |
|---|---|---|
| 5 | Parental predictability | 17: My family had holiday traditions that we did every year (e.g., cooking a special food at a particular time of year, decorating the house the same way). |
| 6 | Physical environment | 36: I lived in a clean house. |
| 6 | Physical environment | 37: I lived in a cluttered house (e.g., piles of stuff everywhere). |
| 6 | Physical environment | 38: In my house, things I needed were often misplaced so that I could not find them. |

### S4. Demographic Analyses

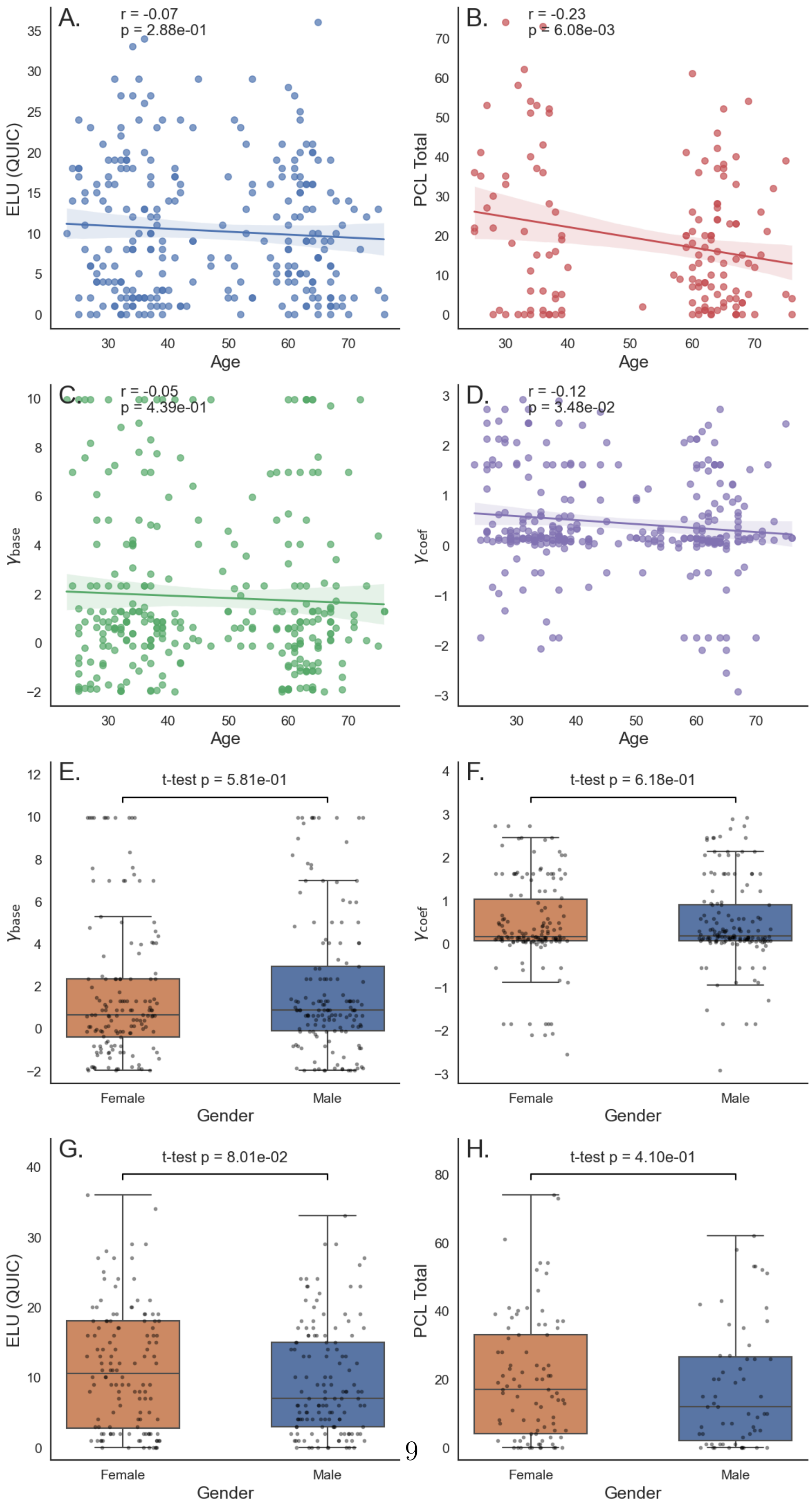

**Figure S4: Demographic effects on behavioral and model parameters.** (A) Relationship between age and early-life unpredictability (QUIC total). (B) Relationship between age and PTSD symptom severity (PCL total). (C) Relationship between age and baseline discounting ($\gamma_{\text{base}}$). (D) Relationship between age and uncertainty sensitivity ($\gamma_{\text{coef}}$). (E) Gender differences in $\gamma_{\text{base}}$. (F) Gender differences in $\gamma_{\text{coef}}$. (G) Gender differences in QUIC total. (H) Gender differences in PCL total.

### S5 Uncertainty is unrelated to trial number

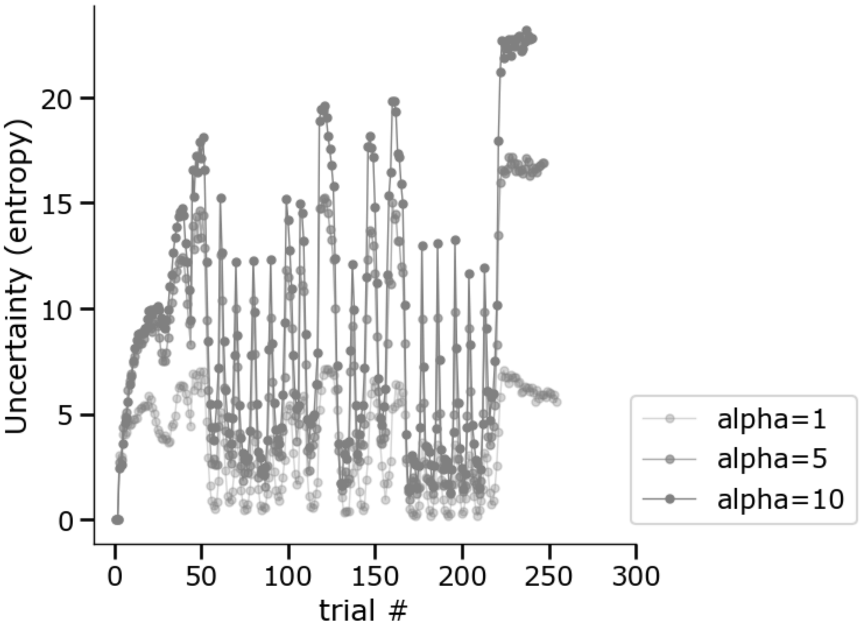


**Figure S5: Entropy/Uncertainty as a function of trials.** Uncertainty and time on task are not correlated in our design, and so any effects of uncertainty-sensitive discounting cannot be attributed to experience within the task.
